# Genomic relationships among diploid and polyploid species of the genus *Ludwigia* L. section *Jussiaea* using a combination of molecular cytogenetic, morphological, and crossing investigations

**DOI:** 10.1101/2023.01.02.522458

**Authors:** D. Barloy, L. Portillo - Lemus, S. A. Krueger-Hadfield, V. Huteau, O. Coriton

**Affiliations:** DECOD (Ecosystem Dynamics and Sustainability), Institut Agro, INRAE, IFREMER 35042 Rennes, France; Molecular Cytogenetics Platform, IGEPP (Institute for Genetics, Environment and Plant Protection), INRAE, Institut Agro, Univ Rennes, 35653, Le Rheu, France; Virginia Institute of Marine Science Eastern Shore Laboratory, Wachapreague, VA, 23480, USA

**Keywords:** GISH, invasive plant, *Ludwigia* L., Onagraceae, polyploidy, phylogenetics

## Abstract

The genus *Ludwigia* L. section *Jussiaea* is composed of a polyploid species complex with 2x, 4x, 6x and 10x ploidy levels, suggesting possible hybrid origins. The aim of the present study is to understand the genomic relationships among diploid and polyploid species in the section Jussiaea. Morphological and cytogenetic observations, controlled crosses, genomic *in situ* hybridization (GISH), and flow cytometry were used to characterize species, ploidy levels, ploidy patterns, and genomic composition across taxa. Genome sizes obtained were in agreement with the diploid, tetraploid, hexaploid, and decaploid ploidy levels. Results of GISH showed that progenitors of *Ludwigia stolonifera* (4x) were *Ludwigia peploides* subsp. *montevidensis* (2x) and *Ludwigia helminthorrhiza* (2x), which also participated for one part (2x) to the *Ludwigia ascendens* genome (4x). *Ludwigia grandiflora* subsp. *hexapetala* (10x) resulted from the hybridization between *L. stolonifera* (4x) and *Ludwigia grandiflora* subsp. *grandiflora* (6x). One progenitor of *L. grandiflora* subsp. *grandiflora* was identified as *L. peploides* (2x). Our results suggest the existence of several processes of hybridization, leading to polyploidy, and possibly allopolyploidy, in the section Jussiaea due to the diversity of ploidy levels. The success of GISH opens up the potential for future studies to identify other missing progenitors in *Ludwigia* L. as well as other taxa.

## Introduction

Polyploidization is widespread in plants and is considered as a major driving force in plant speciation and evolution (Otto & Whitton, 2000; Husband et al., 2013; Alix et al., 2017). Autopolyploid plants arise from the duplication of one genome within one species and allopolyploid plants result from the association of two or more divergent genomes through interspecific hybridization and subsequent genome duplication(Soltis et al., 2015; Alix et al., 2017). Furthermore, some polyploids can arise from both auto-and allopolyploidy events because of their evolutionary histories and are called auto-allo-polyploid. Genomic analyses have revealed that all angiosperms have been subjected to at least one round of polyploidy in their evolutionary history and are thus considered paleopolyploids (Garsmeur et al., 2014). Thus, understanding the origins of polyploid taxa is integral to understanding angiosperm evolution.

Polyploid plants are often thought to be more resilient to extreme environments than diploids because of their increased genetic variation (Husband et al., 2013). Their duplicated genes act as a buffer and can include gene conversion events, activation of transposable elements, chromatin remodelling, and DNA methylation changes (Hollister, 2015). Polyploidy might confer an advantage with both abiotic and biotic stress by increasing tolerance to salt or drought stress or by improving resistance to bioagressors (Van de Peer et al., 2021). Thus, polyploids are able to occupy new ecological niches (Stebbins, 1985; Blaine Marchant et al., 2016) and often show greater adaptability than their progenitors (McIntyre, 2012; Allario et al., 2013; Baniaga et al., 2020; Akiyama et al., 2021).Van de Peer et al. (2021) suggested that as in a constant environment, polyploidization may play an important role in response to habitat disturbance, nutritional stress, physical stress, and climate change (Wei et al., 2019). For example, Baniaga et al. (2020) showed that ecological niches of polyploid plants differentiated often faster than found in their diploid relatives. A polyploid advantage has also been reported in invasive plants and their success in non-native habitats (Te Beest et al., 2012). However, Lobato-de Magalhães et al. (2021) observed little difference in the incidence of each ploidy state within a set of 49 of the world’s most invasive aquatic weeds and concluded there is no consistent evidence of polyploid advantage in invasiveness. Nevertheless, *Spartina anglica*, an invasive neoallopolyploid weed species that appeared around 1890, has increased fitness with its prolific seed production, fertility, and extensive clonal growth as compared to its progenitors (Baumel et al., 2002). A recent study including 50 alien non-invasive aquatic plant species and 68 alien invasive species across various aquatic habitats in the Kashmir Himalayas found that invasive species are largely polyploids whereas non-invasive species tend to diploids (Wani et al., 2018).

*Ludwigia* L., a worldwide wetland genus of 83 species, forms a strongly monophyletic lineage sister to the rest of the Onagraceae. It is currently classified as members of 23 sections (Levin et al., 2003, 2004). Sections were clustered into three main groups by Raven (1963). The first group concerned the Myrtocarpus complex, comprising 14 sections (Raven, 1963; Eyde, 1977; Ramamoorthy, 1979; Zardini & Raven, 1992). The second group included species in the section *Eujussiaea* Munz (Munz, 1942), also referred to as a sect. *Oligospermum* (Raven, 1963) but now correctly called sect. *Jussiaea* (Hoch et al., 1993). The third group combined species in sect. *Isnardia*, sect. *Ludwigia*, sect. *Microcarpium*, and sect. *Miquelia* P.H. Raven (Raven, 1963; Wagner et al., 2007). Liu et al. (2017) provided the first comprehensive molecular phylogeny of *Ludwigia* genus using both nuclear and chloroplast DNA regions. Sixty of 83 species in the *Ludwigia* genus were distributed in the two clades A and B, with the sub-clade B1 which consisted of only sect. *Jussiaea*. This section included seven species: three diploid species (2n=2x=16) (*Ludwigia torulosa* (Arn.) H. Hara, *Ludwigia helminthorrhiza* (Mart.) H. Hara, *Ludwigia peploides* (Kunth) P.H. Raven); two tetraploid species (2n=4x=32) (*Ludwigia adscendens* (L.) H. Hara, *Ludwigia stolonifera* (Guill. &Perr.) P.H. Raven); one hexaploid species (2n=6x=48) (*Ludwigia grandiflora* subsp. *grandiflora*); and one decaploid species (2n=10x=80) (*Ludwigia grandiflora* subsp. *hexapetala*). While most species are native to the New World, particularly South America, two species are restricted to the Old World, *Ludwigia stolonifera* and *Ludwigia adscendens*, in Africa and tropical Asia, respectively (Wagner et al., 2007) (Supplementary Table S1). It is not easy to distinguish between the hexaploid and decaploid species morphologically and both have previously been treated as a single species (*Ludwigia uruguayensis* (Cambess.) H. Hara; (Zardini et al., 1991). Octoploid hybrids between *L. grandiflora* subsp. *hexapetala (Lgh)* and *L. grandiflora* subsp. *grandiflora* (*Lgg*) were found in southern Brazil which for both species is their native area (Zardini et al., 1991). Studies of Liu et al. (2017) confirmed close relationship between *Lgg* and *Lgh*. So, Nesom & Kartesz (2000) suggested that as *Lgg* and *Lgh* shared genomic portions and possible hybridization between them, both species were recognized as subspecies within the single species *L. grandiflora*. However, several authors, including Okada et al. (2009) and Grewell et al. (2016), continue to recognize two distinct species. In this paper, species were named as described by Nesom & Kartesz (2000) and Armitage et al. (2013), i.e., considered as two subspecies of *L*. *grandiflora* (*Lgg* and *Lgh*). So, phylogenetic studies (Liu et al., 2017) revealed that the *L. peploides* (2x) or a relative and the *L. adscendens* (4x) are probably progenitors to *L. stolonifera* and *L. × taiwanensis* (3x), respectively. Furthermore, based on morphological observations, Zardini et al. (1991) suggested that *Lgh* may be result of interspecific hybridization between *Lgg* and *L. hookeri*. So, in view of the diversity of ploidy levels present in the *ludwigia* sect*. Jussiaea*, results of morphological and molecular analysis, polyploid species could be probably the result of hybridization between diploid species or combinations of diploid and polyploid species. In this study, we focused on species belonging to the second group, sect. *Jussiaea*. Most species of the section grow in warm temperate to subtropical moist or wet habitats worldwide. Some of these species, such as *Ludwigia peploides* subsp. *montevidensis* (Kunth) P.H. Raven, *Ludwigia grandiflora* (syn*. L. grandiflora* subsp. *grandiflora*), *Ludwigia hexapetala* (Hook. & Arn.) Zardini, H.Y. Gu & P.H. Raven (syn*. L. grandiflora* subsp. *hexapetala*) (Hook. & Arn.) Zardini, H. Y. Gu & P. H. Raven, can be invasive weeds in wetlands and other wet areas in the USA (Grewell et al., 2016), Europe (Portillo-Lemus et al., 2021), Japan (Hieda et al., 2020), and Korea (Kim et al., 2019). Recently, Méndez Santos and González-Sivilla (2020) revealed that *L. helminthorrhiza* (Mart.) H. Hara must be treated and managed as an invasive alien species in Cuba. Reproductive systems in *Ludwigia* L. are both clonal with production of asexual fragments and sexual with seeds production. Okada et al. (2009) showed that clonal spread through asexual reproduction is the primary regeneration mode of *L. grandiflora* subsp. *grandiflora* and *L. grandiflora* subsp. *hexapetala* in California. Furthermore, Dandelot (2004) reports that all the populations of *L. grandiflora* subsp. *hexapetala* in the French Mediterranean area could have originated from a single clone. Similarly, Reddy et al. (2021) observed low genotypic diversity in both *L. grandiflora* subsp. *grandiflora* and *L. grandiflora* subsp. *hexapetala* in the United State with as example an analysis of multiple invasive populations of *L. grandiflora* subsp. *hexapetala* in Alabama, California, Oregon, Washington, and Florida identified a single genotype.

The aim of this study is to explore the complicated evolutionary history of genus *Ludwigia* L. section *Jussiaea* using a combination of cytogenetic, morphological, and crossing investigations. This is a difficult puzzle to elucidate, with taxa ranging from diploid to decaploid and with both allo-and autopolyploidy involved in the history of these taxa. The occurrence of different ploidy levels of *Ludwigia* species belonging to the same clade might indicate that a diploid species in this clade could be the progenitor of the polyploids analysed. However, while many authors have highlighted the possibility of interspecific hybridization between the species presents in the Jussieae section, there is a lack of data enabling the polyploid origin of these species to be identified, i.e., the auto or allopolyploid origin as well as that of the progenitor species. First, we observed some morphological traits as a simple verification step to prove that the species collected were those expected. Second, we characterized the different species by analysis of their genome size using flow cytometry and their ploidy level by cytogenetic observations. We identified the genomic relationships by Genomic *in situ* Hybridization (GISH) and evaluated the ability of inter-species hybridization after controlled pollination. The genomic relationships between diploid and polyploid species are reported for the first time in sect. *Jussiaea*.

## Material and Methods

### Plant material

Two diploid, two tetraploid, one hexaploid, and one decaploid *Ludwigia* species were analysed. Fifteen plants of *Ludwigia peploides* subsp. *montevidensis* (2x) (hereafter, *Lpm*) and of *L. grandiflora* subsp. *hexapetala* (hereafter, *Lgh*) (10x) were collected in France at the marshes of la Musse (47°14’27.5”N, 1°47’21.3”W) and Mazerolles (47°23’16.3”N, 1°28’09.7”W), respectively. Ten plants of the diploid species *L. helminthorrhiza* (hereafter, *Lh)* was purchased in aquarium store (provider Ruinemans Aquarium B.V. Netherland). Five plants of *Ludwigia adscendens* (L.) H. HARA (4x) (hereafter, *La*), and of *L. stolonifera* (4x) (hereafter, *Ls*) and ten of *L. grandiflora* subsp. *grandiflora* (6x) (hereafter, *Lgg*) were collected in Flores island, Indonesia (Pulau Flores; 8°49’40.8”S, 120°48’39.0”E), Lebanon (Hekr al Dahri; 34°37’54.5”N, 36°01’28.9”E), and the USA (Co. Rd 73, outside Greensboro, AL; 32°61’51.41”N, 87°68’65.4”W), respectively. As all *Ludwigia* species reproduce preferentially by clonal reproduction ; each plant was used as mother plant giving new plants from the development of buds present on its stem which are then used for all experiments (Okada et al., 2009; Glover et al., 2015). The plants were easily maintained in the greenhouse at Institut Agro Rennes - Angers before analysis (Portillo Lemus et al., 2021).

### Morphology

To confirm that the collected *Ludwigia* species corresponded to the expected species, we carried out qualitative observations using simple visual morphological traits such as the colour of the flowers and roots and the pneumatophore form as reported in Supplementary Table S1. Morphological observations for each species were made on at least 30 plants in the greenhouse and confirmed in natura on 15 plants in 15 and 36 populations of *Lpm* and *Lgh* in France, respectively.

### Chromosome counting

At least 40 root tips of 0.5 - 1.5 cm in length were taken for each *Ludwigia* sp. as follows from 15 *Lpm*; ten *Lh*; five *La*; five *Ls*; ten *Lgg* and 15 *Lgh* different plants and were incubated in 0.04% 8-hydroxiquinoline for 2 hours at room temperature in the dark, followed by 2h at 4°C to accumulate metaphases. Chromosome preparations were performed according to procedures detailed in (Książczyk et al., 2011). At least four roots per species were observed. The 4’,6-diamidino-2-phenylindole (DAPI) staining chromosome counts per species were estimated on a total of 20 cells at the mitotic metaphase stage using the visualization software Zen 2 PRO (Carl Zeiss, Germany).

### Genome size estimation by flow cytometry

To explore the genome size among the different *Ludwigia* spp., we used flow cytometry. Approximately 4 mg of fresh roots or leaves from five plants of *Ludwigia* spp. and of fresh leaves from five plants of *Trifolium rupens* (2C DNA = 2.23 pg) *or Zea mays* (2C DNA = 5.55 pg) (Zonneveld, 2019) (used as an internal reference standard for *Lpm*, *Lh* and *Lgh* species and *Ls*, *La*, *Lgg* and *Lgh* species, respectively) were harvested and transferred to a Petri dish. Estimation of genome size for each species was obtained as described by Boutte et al. (2020). For the different *Ludwigia* spp., two or three measures of genome size were made, excepted for *Ls* (only one measure). From each species, the mean ratio of DNA content was calculated (mean + CI (Confidence Interval), p-value= 0.05)). Genome sizes were converted from picograms (pg) to Megabases (Mb) using 1 pg = 978 Mbp (Dolezel et al., 2003).

### Genomic in situ hybridization (GISH)

GISH is used to distinguish chromosomes from different genomes in interspecific/intergeneric hybrids or allopolyploids. Total genomic DNA of a genitor involved in the formation of a hybrid is used at the same time as an unlabeled DNA from another genitor, at a higher concentration, which serves as a blocking DNA, hybridizing with the sequences in common with both genomes. This method is based on repetitive sequences which are more often in plant species-specific. Thus, we compared the level of relatedness between the genomes of the studied species and hypothetical parental species.

DNA was extracted from 30 mg of freeze-dried buds taken from 15 *Lpm*, ten *Lh*, five *Ls*, five *La*, ten *Lgg,* and 15 *Lgh* plants, using the Macherey-Nagel extraction kit NucleoSpin® Food to which we have made following modifications to obtain a polysaccharide free DNA: (1) after lysis step with Buffer CF, we mixed freeze-dried buds with an equivalent volume of PCIA 25:24:1 (parts of phenol, chloroform, isoamyl alcohol) for 5 minutes; (2) then we transferred the whole in a tube containing phase-look gel and centrifuged at 800rpm for 5 minutes (Quantabio, Massachusetts, USA); (3) then the DNA was precipitated using absolute ethanol at -18°C instead of QW and C5 buffers. Finally, the DNA was resuspended after an incubation of 5 min in 100 ml elution buffer with 5 mM TRIS at pH 8.5 at 65°C. 500 ng of total genomic DNA were labelled by random priming with biotin-14-dCTP (Invitrogen by Thermo Fisher Scientific) used as probes.

Total genomic DNA used as a blocking DNA was autoclaved to yield fragments of 100-300 bp. The ratio DNA probe / blocking DNA was 1:50. The hybridized probes correspond to the chromosomes present on the slide (i.e., same species) and genomic DNA (blocking DNA) from different species were used as competitors in to block the common sequences at both species. Genomic In Situ Hybridization (GISH) was carried out as described in Coriton et al. (2019), using a 5 µg of blocking DNA (∼50-fold excess). Biotinylated probes were immunodetected by Texas Red avidin DCS (Vector Laboratories, Burlingame, CA, USA) and the signal was amplified with biotinylated anti-avidin D (Vector Laboratories). The chromosomes were mounted and counterstained in Vectashield (Vector Laboratories) containing 2.5µg/mL 4’,6-diamidino-2-phenylindole (DAPI). Fluorescence images were captured using an ORCA-Flash4 (Hamamatsu, Japan) on an Axioplan 2 microscope (Zeiss, Oberkochen, Germany) and analysed using Zen 2 PRO software (Zeiss, Oberkochen, Germany). For each *Ludwigia* species, at least three independent slides were made with a total of 20 cells observed per species. The images were processed using Photoshop v.8.0.1 (Adobe Systems Inc., San Jose, CA, USA).

### Controlled interspecific crosses

Controlled interspecific pollinations were carried out in the greenhouse between *Ludwigia* species which putatively shared the same parental genome. Thus, interspecific hybridizations were made between *L. peploides* subsp. *montevidensis*, *L. stolonifera* and/or *L. grandiflora* subsp. *hexapetala* used as male or as female. Ten plants of each species were used for crosses. *Ludwigia* spp. produced flowers on a shoot until July to October, with at one time only one flower per shoot at the good stage of mature for pollination. To carry out interspecific pollinations, flowers were enclosed in cellophane bags to protect them from external pollen before and after pollination. Flowers used as ‘female’ were emasculated before anthesis. A mix of pollen from flowers of five different plants for each of other species was used to pollinate emasculated flowers. Between two to 25 interspecific crosses were made according to the availability of flowers. To control efficiency of pollination in greenhouse, we also conducted at the same time 45, 75 and 50 intraspecific crosses for *Lpm, Lgh* and *Ls*, respectively.

Pollination success for interspecific crosses was estimated by the number of fruits, fruit size and weight, the number of seeds, viable plantlets, and the number of plants ultimately produced. For intraspecific crosses, the number of fruits obtained were noted.

## Results

### Morphological traits of Ludwigia species

The qualitative traits observed in the species collected, namely the color of flowers and roots and the pneumatophore form, were consistent with those reported for these species in literature (see Supplementary Table S1 for traits and authors).

For the diploid species, red roots, yellow flowers, and rare cylindric pneumatophores were observed in *Lpm*. In contrast, in *Lh*, we observed red roots, creamy white petals with narrow yellow base, and abundant, clustered conical pneumatophores (Figure 1). For the tetraploid species, *La* had pink roots, white petals with yellow base, and had few conical pneumatophores. *Ls* had white roots, petal color light yellow and similar form of pneumatophores as those of *La*. For the hexaploid species *Lgg*, only roots were observed and were pink. The decaploid species *Lgh* had white roots, flowers with yellow petals, and few, long cylindrical pneumatophores per node. Color of roots and pneumatophore number and form were confirmed in natura for the different populations of *Lpm* and *Lgh* observed (Figure 1).

**Figure 1.**
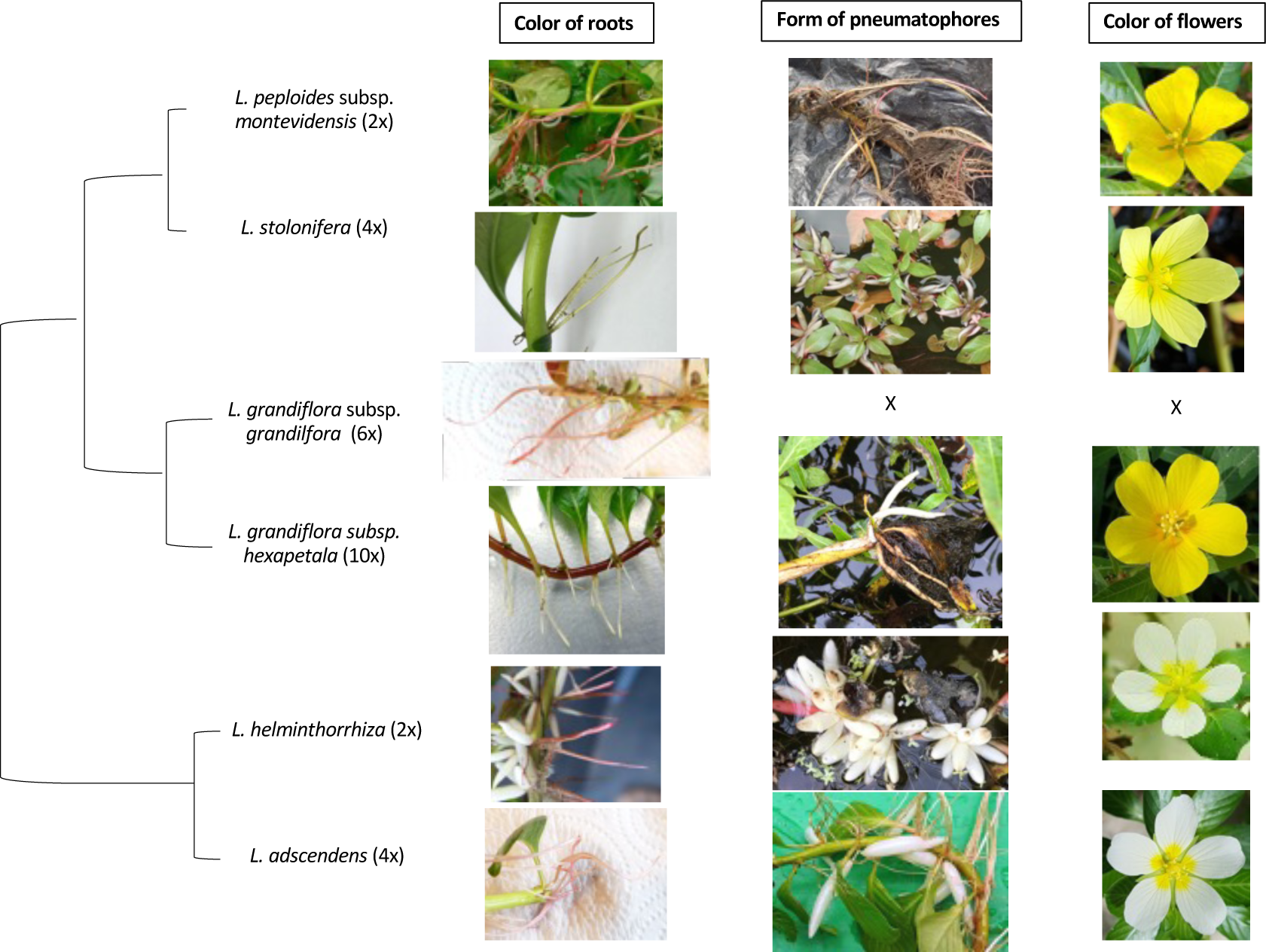
Morphological traits of *Ludwigia* L. species in section *Jussiaea*. *Ludwigia* L. species are classified in a phylogenic tree as proposed by Liu et al. (2017). Three morphological traits were observed (color of roots, pneumatophore form, color of flower).

### Genome size and ploidy level

The chromosome numbers were as expected: for both diploids, *Lpm* and *Lh,* 2n = 16; for both tetraploids *Ls* and *La,* 2n =32; for hexaploid *Lgg*, 2n=48 and for decaploid *Lgh,* 2n=80 (Table 1, Supplementary Figure S1). *Ludwigia* spp. exhibited an ∼0.77-fold range of C-values. The lowest value, 0.53 pg/2C, was found in *Lpm* and the highest, 2.9pg/2C, in *Lgh* (Table 1, Supplementary Figure S2). The tetraploid species *Ls* (1.07pg/2C) and *La* (1.06pg/2C) have C-values that were twice that the value for the diploid *Lpm* (0.53pg/2C) and *Lh* (0.55pg/2C). The hexaploid species *Lgg* had C-value 1.77pg/2C. Thus, the genome size by ploidy level revealed that the monoploid genome sizes (1Cx-value, 0.133-0.147 pg) of the tetraploid, hexaploid, and decaploid species are the same (0.34-0.49 pg/1Cx). The difference is accounted for by the higher ploidy levels.

**Table 1:** Ploidy levels, chromosome numbers and genome sizes estimated by flow cytometry in *Ludwigia* L. spp. sect. *Jussiaea*. Species names are mentioned according to the revised nomenclature by Hoch et al. (2015). Genome sizes were converted from picograms (pg) to Megabases (Mb) using 1 pg = 978 Mbp.

| Species name | Ploidy and chromosome numbers | DNA nuclear content (1C in pg) | Genome size (Mb) |
| --- | --- | --- | --- |
| *Ludwigia peploides* subsp. *montevidensis* (Lpm) | 2n= 2x= 16 | 0.265 | 261.7 +/- 9.2 |
| *Ludwigia helminthorrhiza* (Lh) | 2n= 2x= 16 | 0.275 | 267.7 +/- 1.7 |
| *Ludwigia adscendens* (La) | 2n= 4x= 32 | 0.53 | 498.6 +/- 31.4 |
| *Ludwigia stolonifera* (Ls) | 2n= 4x= 32 | 0.535 | 522 |
| *Ludwigia grandiflora* subsp. *grandiflora* (Lgg) | 2n= 6x= 48 | 0.885 | 863.9 +/- 2.6 |
| *Ludwigia grandiflora* subsp. *hexapetala* (Lgh) | 2n= 10x= 80 | 1.045 | 1418.9 +/- 107.3 |

*Ludwigia* genome sizes of diploid and tetraploid species were similar between species with the same ploidy level and varied proportionally with ploidy levels (i.e., 2x:::260 Mb, 4x::: 500 Mb; Table 1, Supplementary Figure S2). The genome size of hexaploid and decaploid species were closer than those expected with regard to ploidy level (i.e., ratio (6x/2x) = 1.07; ratio (10x/2x) = 1.06; Table 1) with 864 Mb and 1419 Mb, respectively.

### Genomic relationships using the GISH technique

For the diploid species, when we hybridized slides of *Lpm* with a *Lpm* probe (red) and *Lh* blocking DNA (grey), 16 chromosomes were tagged in red signals and zero chromosome showed a grey signal (Figure 2A). Thus, the *Lh* blocking DNA did not block any sequence present in the *Lpm* probe, meaning that no *Lh* genome was shared with *Lpm*. But, when slides of *Lh* were hybridized with a *Lh* probe and *Lpm* blocking DNA, ten chromosomes of *Lh* showed grey signal corresponding to *Lpm* chromosomes (Figure 2B). This observation suggests genome homology with the *Lpm* genome but four chromosomes were stained in red, meaning that there are nevertheless differences in *Lpm* and *Lh* genomes. Due to the absence of chromosomes marked by *Lh* blocking DNA in *Lpm*, we can suggest that even some homology exist, *Lpm* and *Lh* most likely correspond to different genomes, arbitrarily noted A for *Lpm* and B for *Lh*.

**Figure 2.**
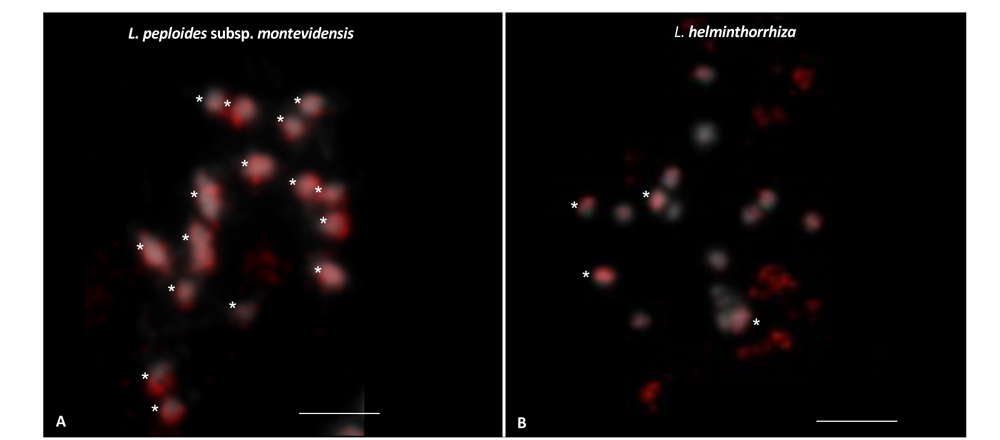
Genomic in situ hybridization (GISH) on mitotic metaphase chromosomes from *Ludwigia peploides* subsp. *montevidensis* (2n= 2x =16) using *Ludwigia peploides* subsp. *montevidensis* probe (2x) (red) and *Ludwigia helminthorrhiza* (2x) (10µg) as blocking DNA (A) and from *L. helminthorrhiza* (2n= 2x =16) using *L. helminthorrhiza* probe (2x) and *L. peploides* subsp. *montevidensis* (2x) (10µg) as blocking DNA (B). Thus, GISH reveals specifically 16 red signals (white stars) and 0 *L. peploides* subsp. *montevidensis* chromosomes (grey) (A) and 4 red signals (white stars) and 10 *L. helminthorrhiza* chromosomes (grey) (B). Chromosomes were counterstained with DAPI (grey). Bar represents 5 µm.

For the tetraploid species *Ls* and *La*, we hybridized *Ls* slides with a *Ls* probe and three different blocking DNA combinations from species having different ploidy levels – *Lpm* (2x), *Lh* (2x) and *La* (4x) – and for *La* slides, with a *La* probe and *Lh* blocking DNA (Table 2, Figure 3). When *Lpm* DNA was hybridized over *Ls*, the blocking DNA *Lpm* blocked 16 chromosomes (grey) and the other 16 chromosomes tagged in red by the *Ls* probe (Figure 3A). A similar result was obtained with the blocking DNA of *Lh*, with 16 chromosomes showing red signals and 16 grey (Figure 3B). Thus, the tetraploid *Ls* would be the result of an interspecific hybridization between the two diploid species *Lpm* and *Lh.* Based on the genome naming proposed here, the genomic composition of *L. stolonifera* could be AABB.

**Figure 3:**
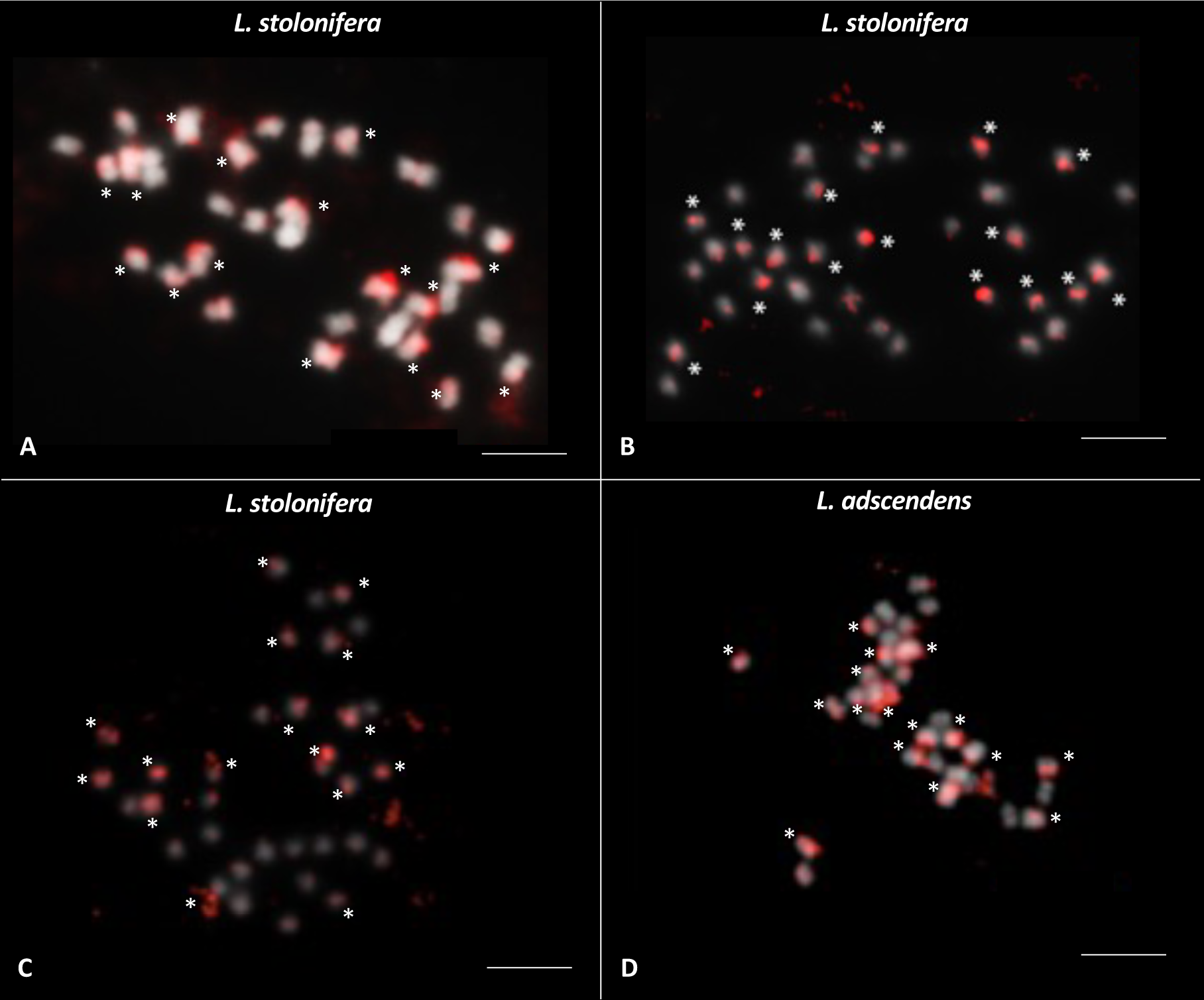
Genomic in situ hybridization (GISH) on mitotic metaphase chromosomes from the tetraploid species, *Ludwigia stolonifera* and *Ludwigia adscendens* (2n= 4x =32). GISH was carried out for *L. stolonifera* using *L. stolonifera* probe (4x) (red) and *Ludwigia peploides* subsp. *montevidensis* (2x) (10µg) as DNA blocking (A), *Ludwigia helminthorrhiza* (2x) as block (B) and *L. adscendens* (4x) as block (C) and for *L. adscendens* (4x) using *L. adscendens* probe (4x) (red) and *L. helminthorrhiza* (2X) (10µg) as block (D). Thus, GISH revealed for *L. stolonifera* specifically 16 red signals (white stars) and 16 *L. peploides* subsp. *montevidensis* chromosomes (grey) (A), 16 red signals (white stars) and 16 *L. helminthorrhiza* chromosomes (grey) (B), 16 red signals (white stars) and 16 *L. adscendens* chromosomes (grey) (C) and for *L. adscendens* 16 red signals (white stars) and 16 *L. helminthorrhiza* chromosomes (grey) (D). Chromosomes were counterstained with DAPI (grey). Bar represents 5 µm.

**Table 2:** Results of GISH with different *Ludwigia* L. probes (red) combined with blocking DNA (grey) on *L. peploides* subsp. *montevidensis* (*Lpm*), *L. helminthorrhiza* (*Lh*), *L. adscendens* (*La*), *L. grandiflora* subsp. *grandiflora* (*Lgg*) and *L. grandiflora* subsp. *hexapetala* (*Lgh*) chromosomes. Chromosomes of one species tagged in red correspond to DNA of this species and chromosomes tagged in grey are blocked by DNA of others species.

| Blocking DNA \ Slide of chromosomes | *Lpm*<br>(2n = 16) | *Lh*<br>(2n = 16) | *Ls*<br>(2n = 32) | *La*<br>(2n = 32) | *Lgg*<br>(2n = 48) | *Lgh*<br>(2n = 80) |
| --- | --- | --- | --- | --- | --- | --- |
| *Lpm* (2n = 16) |  | 4 red signals<br>+ 10 grey signals | 16 red signals + 16 grey signals |  | 32 red signals + 16 grey signals | 48 red signals + 32 grey signals |
| *Lh* (2n = 16) | 16 red signals |  | 16 red signals + 16 grey signals | 16 red signals + 16 grey signals | 48 red signals (8 less intense) | 80 red signals (16 less intense) |
| *Ls* (2n = 32) |  |  |  |  | 32 red signals + 16 grey signals | 48 red signals + 32 grey signals |
| *La* (2n = 32) |  |  | 16 red signals + 16 grey signals |  | 48 red signals (16 more intense) | 80 red signals (16 less intense) |
| *Lgg* (2n = 48) |  |  |  |  |  | 32 red signals + 48 grey signals |

After use of *La* blocking DNA over *Ls* chromosomes, we observed 16 chromosomes tagged in red and 16 chromosomes tagged in grey (Figure 3C). The hybridization performed with *Lh* blocking DNA on the second tetraploid, *La*, identified 16 red chromosomes and 16 grey chromosomes (Figure 3D). Both results suggested that the two tetraploid species *La* and *Ls* shared a same genome coming from *Lh* (BB component). Thus, *Lh* would also be one of the components of the tetraploid *La*, with a XXBB putative genome composition, where the XX genome corresponds to an unknown *Ludwigia* diploid species.

For the hexaploid species *Lgg*, slides of *Lgg* were hybridized with a *Lgg* probe and four blocking DNA of different ploidy levels – *Lpm* (2x), *Lh* (2x), *Ls* (4x), *La* (4x), and *Lgh* (10x) Table 2). The *Lpm* competitor DNA blocked 16 chromosomes (tagged in grey) and 32 chromosomes showing red signals were hybridized with the *Lgg* probe DNA (Figure 4 A). A similar hybridization was obtained with the *Ls* blocking DNA in which slides of *Lgg* had 16 grey chromosomes and 32 chromosomes with red signals (Figure 4 B). Thus, the hexaploid species *Lgg* contains an identical genomic component found in *Ls* (4x) and in *Lpm* (2x; i.e., AA genomic part).

**Figure 4:**
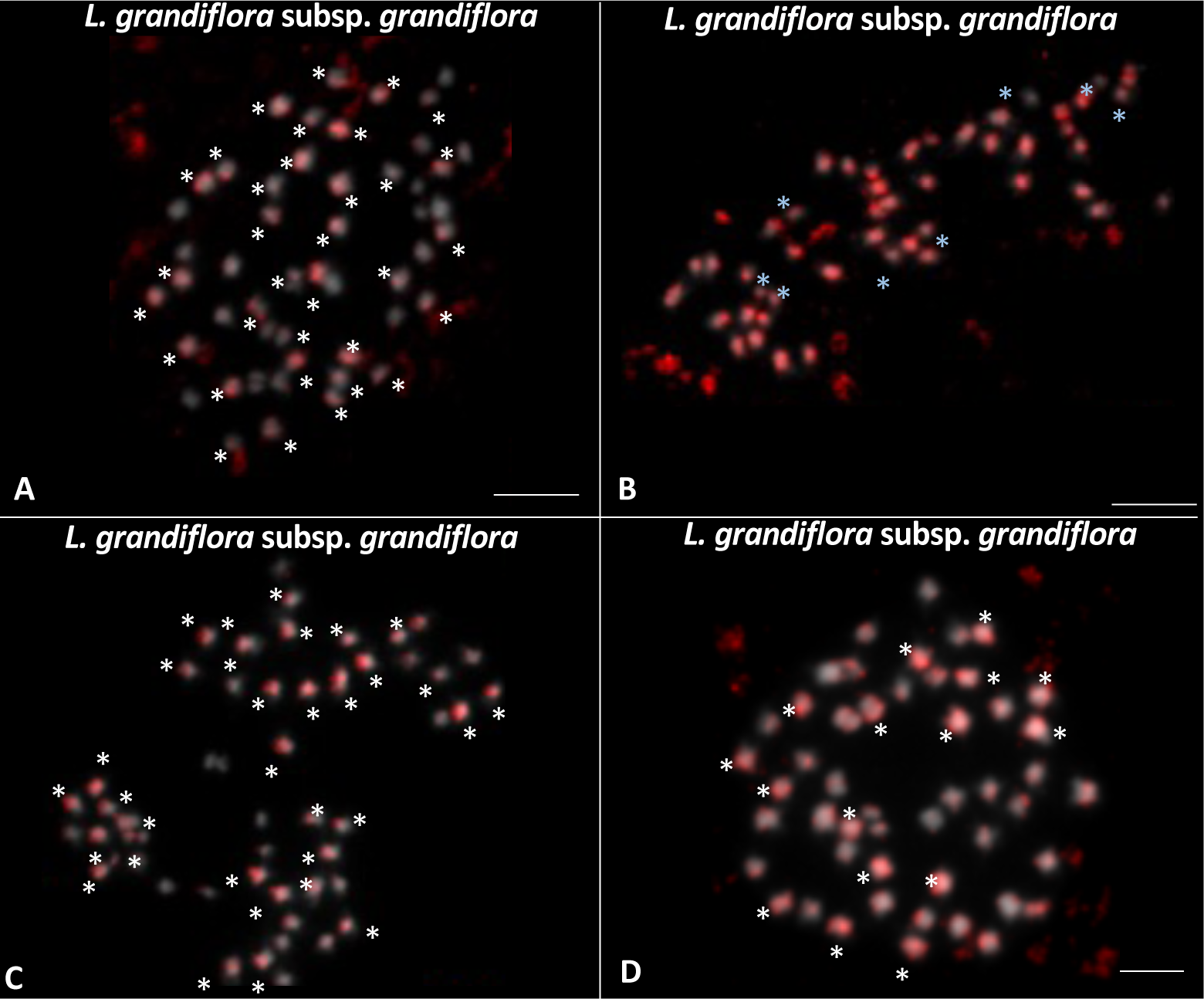
In situ genomic hybridization analyses of somatic metaphase chromosomes from *Ludwigia grandiflora subsp. grandiflora* (2n=6x=48) using *L. grandiflora subsp. grandiflora* probe (6x) (red) and *Ludwigia peploides* subsp. *montevidensis* (2x) (**A**), *Ludwigia helminthorriza* (2x) (10µg) as block (**B**), *Ludwigia stolonifera* (4x) (10µg) as block (10µg) as block (**C**), *Ludwigia adscendens* (4x) (10µg) as block (**D**), *Ludwigia grandiflora* subsp. *hexapetala* (10x) as block (**E**). Thus, GISH reveals specifically 32 red signals (white star) and 16 *L. peploides* subsp. *montevidensis* chromosomes (grey) (**A**), 48 red signals with 8 present less intensity (white star) (**B**), 32 red signals (white star) and 16 *L. stolonifera* chromosomes (grey) (**C**) and 48 red signals with 16 present more intensity (white star) (**D**). Chromosomes were counterstained with DAPI (grey). Bar represents 5 µm.

Hybridizations performed on slides of *Lgg* with *Lh* (2x) and *La* (4x) blocking DNA exhibited hybridization profiles that were more challenging to interpret with 48 red chromosomes, but with different hybridization intensities (with 16 more intense signals with *La* blocking DNA and 8 less intense signals with *Lh* blocking DNA (Table 2, Figures 4C, 4D). The 16 more intense signals could correspond to a 2x component (16 chromosomes) specific to *Lgg*.

For the decaploid species, *Lgh*, slides were hybridized with a *Lgh* probe and five blocking DNA of different ploidy levels, including *Lpm*, *Lh*, *Ls*, *La* and *Lgg*, respectively (Table 2). The *Lpm* DNA competitor blocked 32 chromosomes with grey signals whereby 48 chromosomes showing red signals (Figure 5A). An identical hybridization result was obtained with the *Ls* blocking DNA with 48 chromosomes with red signals and 32 grey chromosomes (Figure 5C). Thus, the 2x component, *Lp*, also present in *Ls* (4x), is found in a double dose (32 chromosomes) in *Lgh* (10x). The results obtained with the *Lh* and *La* DNA blocking showed 80 red chromosomes but 16 with lower intensity (Table 2, Figures 5B, 5D). After GISH hybridization of *Lgg* (6x) DNA on *Lgh* (10x) chromosomes, 32 of 80 *Lgh* chromosomes showed a red signal (Figure 5E). This result revealed that *Lgg* was probably one of progenitors of *Lgh*.

**Figure 5:**
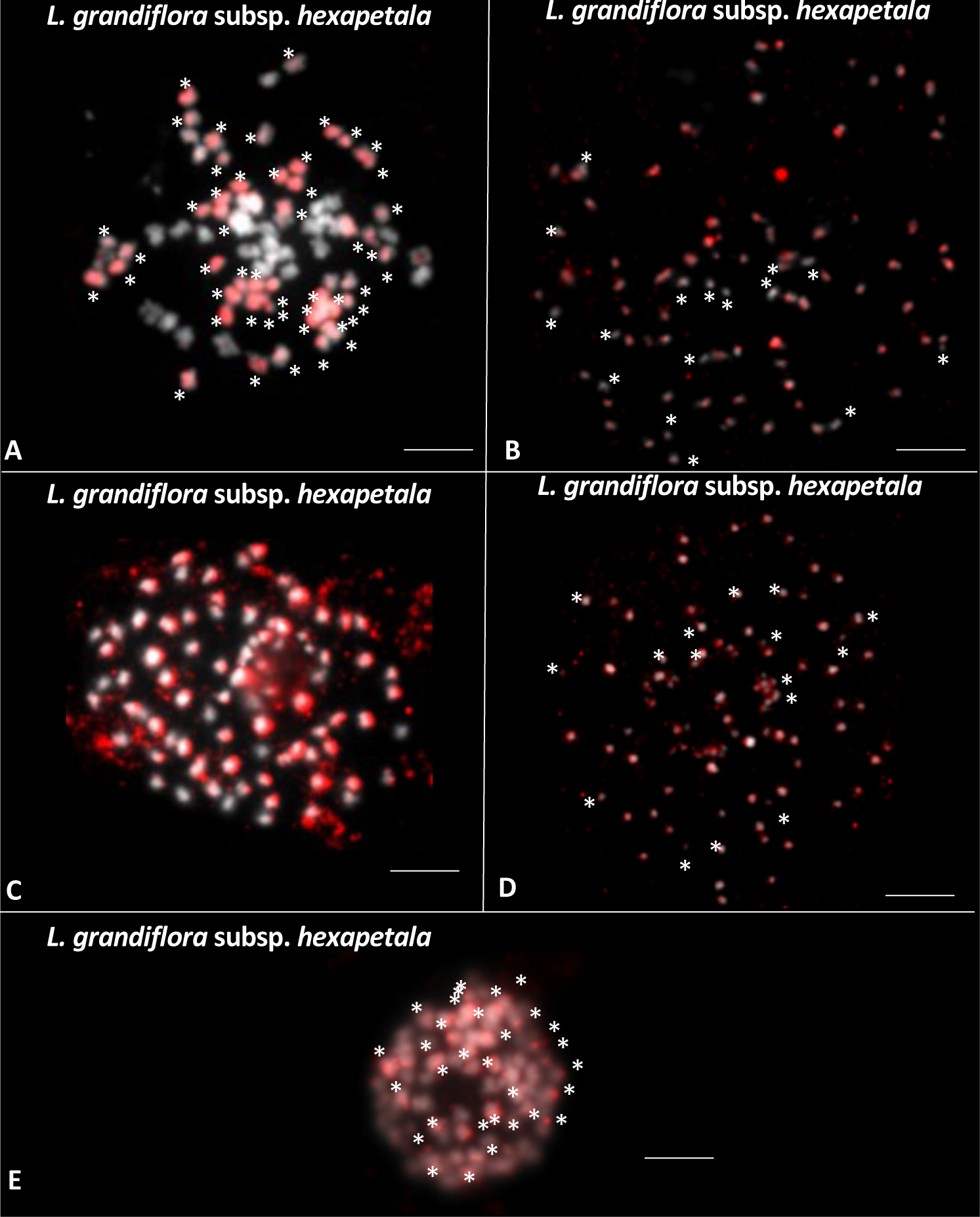
In situ genomic hybridization analyses of somatic metaphase chromosomes from *Ludwigia grandiflora* subsp. *hexapetala* (2n=10X=80) using *L. grandiflora* subsp. *hexapetala* probe (10x) (red) and *Ludwigia peploides* subsp. *montevidensis* (2x) (10µg) as block (**A**), *Ludwigia helminthorrhiza* (2x) as block (**B**), *Ludwigia stolonifera* (4x) (10µg) as block (**C**), *L. adscendens* (4x) as block (**D**) and *Ludwigia grandiflora* subsp. *grandiflora* (6x) as block (**E**). Thus, GISH reveals specifically 48 red signals and 32 *L. peploides* subsp. *montevidensis* chromosomes (grey) (**A**), 80 red signals and 16 present less intensity (white stars) (**B**), 48 red signals and 32 *L. stolonifera* chromosomes (grey) (**C**), 80 red signals and 16 present less intensity (white stars) (**D**) and 32 red signals and 48 *L. grandiflora* subsp. *grandiflora* (grey) (E). Chromosomes were counterstained with DAPI (grey). Bar represents 5 µm.

### Interspecific hybridization

Interspecific hybridization between species sharing the AA genome were carried out and reproductive success was observed by fruit production when the species used as female possessed the lower ploidy level (Figure 6, Supplementary Table S2). No fruits were obtained after crosses between *Ls* (4x) used as female and *Lp* (2x) used as male or between *Lgh* (10x) used as female and *Lpm* (2x) or *Ls* (4x) used as male. Thus, all interspecific crosses with the diploid species *Lpm* (2x) used as female and *Ls* (2x) or *Lgh* (10x) used as male gave fruits showing similar weight and length (Figure 6, Supplementary Table S2). The fruits obtained from the *Lpm* (2x) x *Lgh* (10x) crosses had very large seeds whose development led to the bursting of the fruit walls (Supplementary Figure S3). However, only 53.4% and 3.9% of seeds from *Lpm* (2x) x *Ls* (4x) and *Lpm* (2x) x *Lgh* (10x) crosses germinated. If all germinated seeds gave plantlets for *Lpm* (2x) x *Ls* (2x) crosses, only three plants developed for *Lpm* (2x) x *Lgh* (10x). Finally, no plants survived 90 days after seedling, as all plants showed chlorotic signs and at the end of the observation period, they were not able to survive (Figure 6, Supplementary Table S2, Supplementary Figure S3). Similarly, fruits were produced after *Ls* (4x) x *Lgh* (10x) crosses with a mean number of seeds per fruit of 23.5 (Figure 6, Supplementary Table S2) but no seed has germinated. Unfortunately, chlorotic plants from *Lpm* (2x) x *Ls* (4x) and *Lpm* (2x) x *Lgh* (10x) crosses did not develop enough roots for chromosome observations. For control intraspecific crosses *Lpm* x *Lpm*, *Lgh* x *Lgh* and *Ls* x *Ls*, all crosses produced fruits revealing effectiveness of the greenhouse pollination conditions.

**Figure 6:**
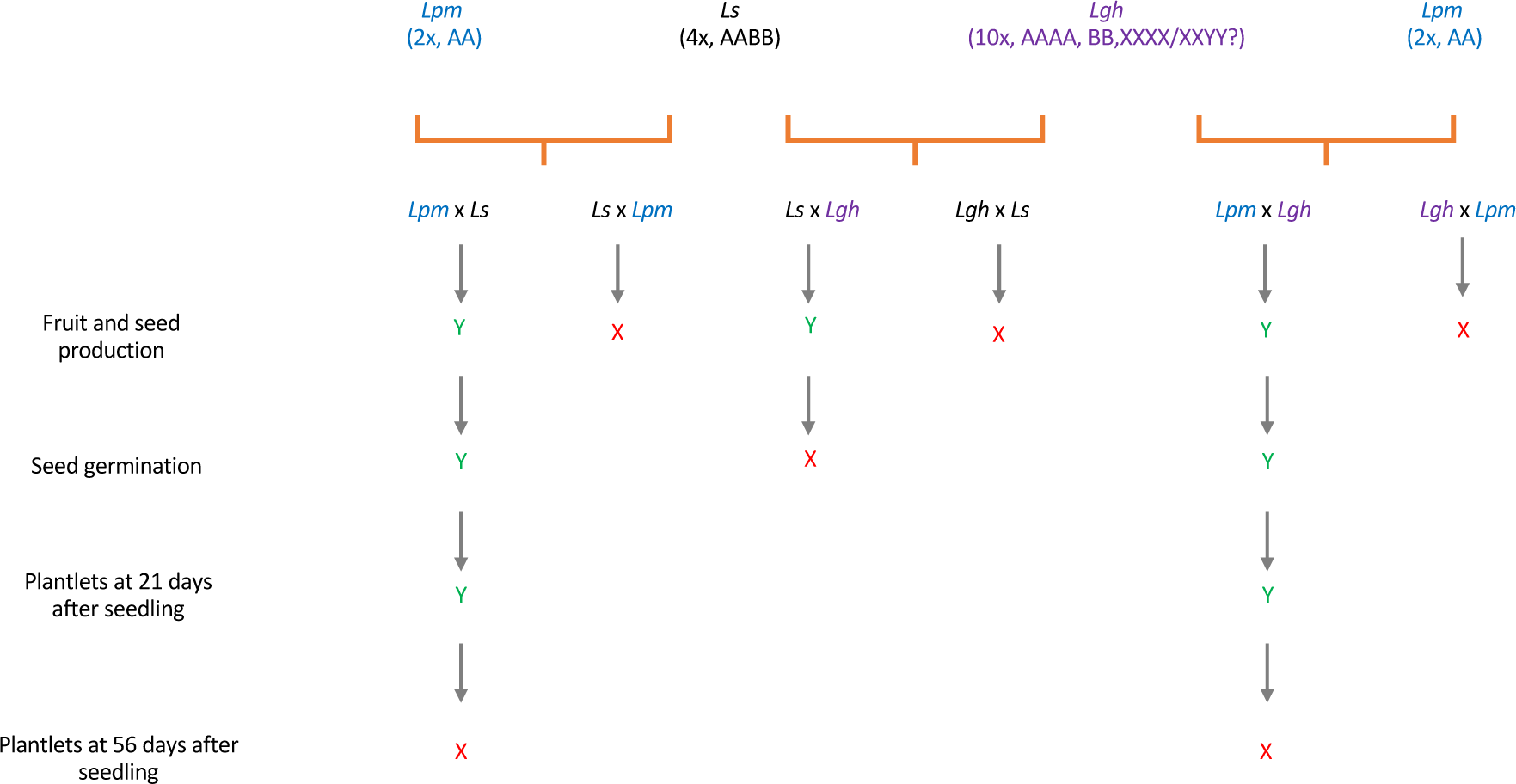
Reproductive success after controlled interspecific crosses between different Ludwigia sp belonging to the section Jussiaea. Interspecific hybridizations were made between the three species, *Ludwigia peploides* subsp *montevidensis* (*Lpm)*, *Ludwigia stolonifera* (*Ls*) and/or *Ludwigia grandiflora* subsp. *hexapetala* (*Lgh*) used as female or male. All species possess same genome A: *Lpm* (2x, AA); *Ls* (4x, AABB); *Lgh* (10x, AAAA BB XXXX or XXYY). Y letter (green) indicated success of step and the cross (red) failed of the corresponding step. Intraspecific crosses (*Lgh* x *Lgh*; *Lpm* x *Lpm*; *Ls* x *Ls*) were used to control pollination efficiency in greenhouse and all pollinated flowers produced fruits.

## Discussion

To better understand the evolutionary history of genus Luwigia, we have evaluated the genomic relationships between diploid and polyploid species using the molecular cytogenetic and crossing investigations.

### Validation of Ludwigia species sect. Jussieae studied and identification of new discriminating traits

Wagner et al. (2007) summarized the complex history of the Onagraceae. The genus *Ludwigia* forms a lineage separate from the rest of the Onagraceae family (Eyde, 1981, 1982) The long-standing taxonomic confusion surrounding aquatic *Ludwigia* species required a approach combining morphometric and cytogenetic evaluations to differentiate the species and improve taxonomic identification (Grewell et al., 2016). Furthermore, distinguishing *Ludwigia* species represents a real challenge.

In this study, qualitative morphological traits were observed for the six Lg ssp. grown in a common garden, allowing compare these species growing under the same conditions. Our results confirmed that all plants collected corresponded to the expected species. However, our cross observations of the different species in a common garden revealed additional differences between these species. For example, the red roots of *Lpm* were never described before, but are visible on the seedlings as soon as the seeds germinate until the plant reaches maturity in natura (Supplementary Figure S4). *Lh* plants studied had these same characteristics as those described (Rocha & Melo, 2020), but the petals were more creamy-white than white and were sharply narrow at the petiole. We found that the pneumatophore form, petal and root coloration could also differentiate these species (Figure 1). For the tetraploid species, flowers of *La* are described as creamy white petals with yellow at the base (Wagner et al., 2007) but we observed white petals similar to *Lh* (Figure 1). As *Ls* had light yellow petals, the flower color may be a good characteristic with which to distinguish these two tetraploid species in natura (Supplementary Figure S4). For the hexaploid species *Lgg*, we only saw pink roots and more morphological investigations are required. Finally, the decaploid species *Lgh* had white roots and bright yellow petals (Figure 1).

Grewell et al. (2016) reported that distinguish in field *Lgg* and *Lgh* was complicated. Nesom & Kartesz (2000) suggested that few morphological distinctions between *Lgh* and *Lgg* exist and broadly overlapping: plants with larger leaves and flowers and less dense vestiture characterize *Lgh*, whereas smaller leaves and flowers and denser vestiture would describe *Lgg*. However, comparing flower morphology in sterile and fertile French *Lgh* populations, two flower sizes were observed which may call into question the criterion for distinguishing flower size between *Lgh* and *Lgg* (Supplementary Figure S4, (Portillo Lemus et al., 2021).

As for the distinction between *Lpm* and *Lgh*, the differences in stipule shape are often cited, reniform for *Lpm* and oblong and acuminate for *Lgh* (Thouvenot et al., 2013), but this character is also not easily used. For all these reasons, we propose new criteria to help field managers based on the root color. *Lpm* has red roots, whereas *Lgh* has white roots. Importantly, this character can be observed at different stages of plant development (Supplementary Figure S4). *Lgg* seems to have pink roots at a young plant stage. Whether this characteristic is also true at all stages of *Lgg* development, it could also be a promising way to distinguish *Lgg* and *Lgh*.

### Genomic relationships and origins of polyploids in section Jussieae

We propose the first hypotheses regarding diploid-polyploid relationships of *Ludwigia* diploid to decaploid species belonging to the section *Jussiaea* (Figure 7). The diploid species studied here were composed of two different genomes, we have called AA and BB for *Lpm* and *Lh*, respectively. Both diploid *Lpm* and *Lh* were the progenitors of *Ls,* with the latter composed of AABB (Figure 3). We also found that *Lh* was a progenitor of *La* (BB), sharing same genome with *Ls* even though the *La*, native to Asian-Pacific, and *Ls*, native to African, do not currently co-occur (Supplementary Table S1). Our results are in agreement with the phylogenetic analysis of Liu et al. (2017) who suggested through analysis of nuclear DNA sequences that *Lp* or a close relative contributed to the origin of *Ls* and shared a same genome (here designated as genome AA). Similarly, Liu et al. (2017) reported that *L. adscendens* (4x) is close to *L. helminthorrhiza* (2x) (genome BB). GIS analysis revealed that *Lh* and *Ls* shared at least one genome, which was not shown by Liu et al (2017) phylogenetic analysis.

**Figure 7:**
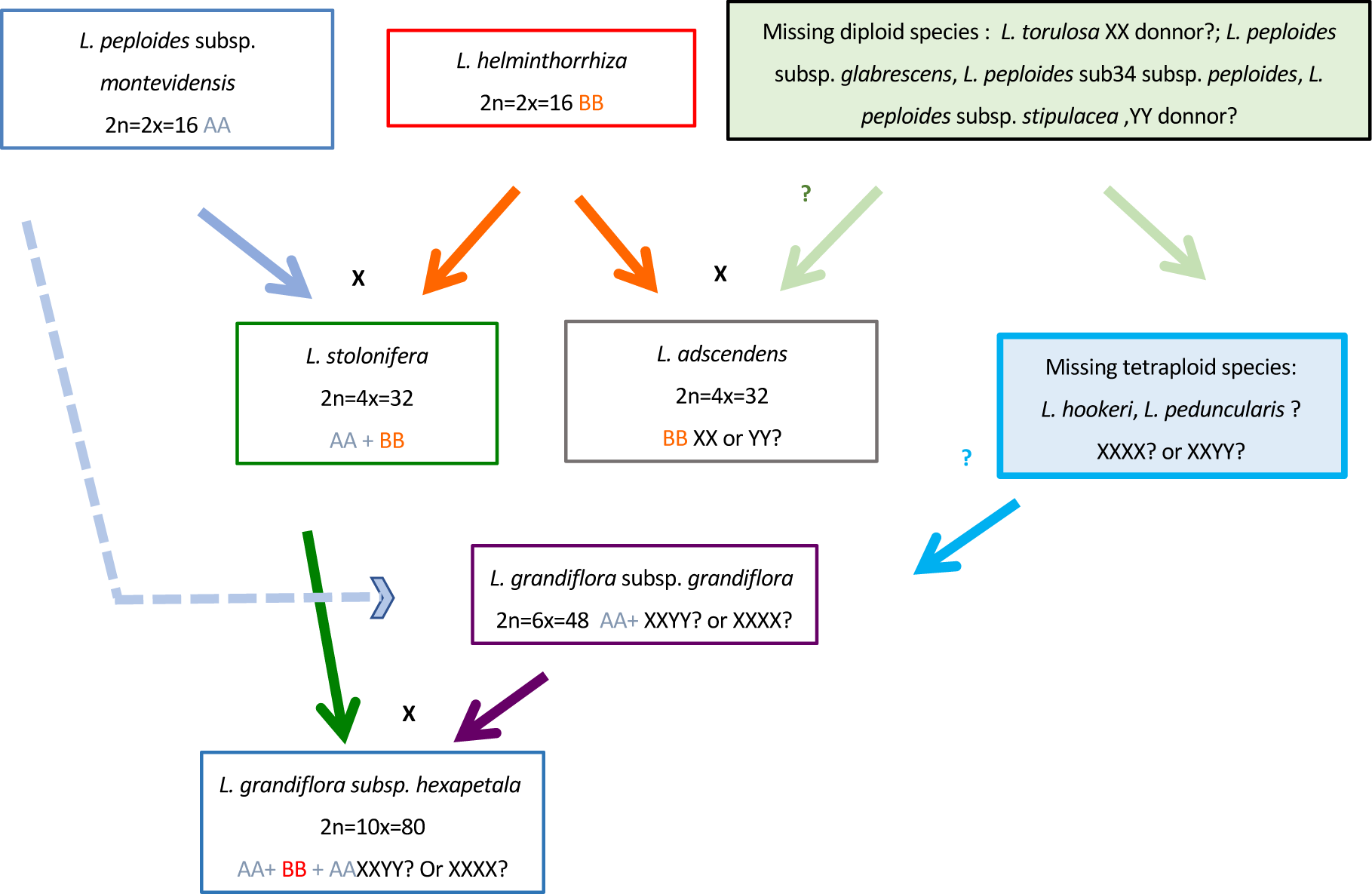
Hypothetical phylogenetic history of *ludwigia* species of section Jussieae

Furthermore, considering the genome sizes of both diploid species *Lpm* and *Lh* and assuming additivity, our genome size data fit perfectly with our scenarios of tetraploid *Ls* and *La* origin. On the other hand, we showed that *Lpm* also participated for one part (2x) to the origin of the hexaploid *Lgg* genome. The decaploid species *Lgh* seems to have emerged from interspecific hybridization and allopolyploidization events between the tetraploid species *Ls* (4x) and the hexaploid species *Lgg*. (Liu et al., 2017) also demonstrated a close relationship between *Lgg* and *Lgh* using nuclear and chloroplast DNA regions as molecular markers. In addition, *Lgh* shares the same pneumatophore form as *Lpm* and the same root color as *Ls*, which may provide further evidence that both species are progenitors of *Lgh*.

All chromosomes of *Lgg* and *Lgh* were tagged by *Lh* blocking DNA, but had strong or light hybridization intensities for 16 chromosomes respectively. The intensity of fluorescence could be explained by repetitive sequences shared among closely related species or specific for given species. Thus, Liu et al. (2008) could distinguish the subgenomes of Triticeae allopolyploids due to differences in element abundance and the resulting probe signal intensity. In addition, in a Silene hybrid, Markova et al. (2007) showed that the intensity of fluorescence varied quantitatively based on the relatedness of the species. These results may suggest genome divergence between *Lgg* or *Lgh* and *Lh*. The intensity level of the signal over the majority of the chromosomes likely indicates a mixing of genomic sequences between parental genomes, in particular for the *Lh* genome (BB), in the hexaploid and decaploid formation. The effectiveness of GISH is much reduced, with clear evidence of considerable mixing of genomic sequence between parental DNA. Lim et al. (2007) have shown that within 1 million years of allopolyploid *Nicotiana* divergence, there is considerable exchange of repeats between parental chromosome sets. After *c.*5 million years of divergence GISH fails. Repetitive sequences, including dispersed repeats, such as transposable elements (Tes), or tandem repeats such as satellite DNAs, represent an important fraction of plant genomes that impact evolutionary dynamics (Vicient & Casacuberta, 2017; Giraud et al., 2021). Yet, no exhaustive investigations have been undertaken to evaluate the nature and dynamics of repetitive sequences between different species of *Ludwigia* that probably diversified since hexaploid and decaploid events when the *Ludwigia* family originated at least 50 m.y. ago (Raven & Tai, 1979).

### Success of interspecific hybridization and contribution to origin of Ludwigia species, sect. Jussieae

In addition to these results, interspecific crosses between *Ludwigia* species sharing the A genome produced fruits only when the female parent possessed lower ploidy level suggesting that efficiency of pollination was possible through the presence of the same genome in both species. In interspecific crosses differences also exist according to the ploidy level of the female parent. For example, in *Brassica ssp.*, more hybrids formed when allotetraploid species, *Brassica napus* is used as female in crosses with diploid species used as male (Kerlan et al., 1992). In contrast, several crosses between *Triticum aestivum* L. and diploid wild relatives were successful when the female parent had the lower chromosome number (Sharma, 1995). Liu et al. (2017) observed through the cp tree analysis that *La* and *Ls* are grouped suggesting that both decaploid species shared at least one maternally inherited genome, probably the BB genome from *Lh*. Unfortunately, *Lh* was not include in cp tree analysis by Liu et al. (2017). The combined data from the interspecific crosses carried out in this study and the phylogenetic analysis carried out by Liu et al. (2017) allows us to hypothesize that in *Ludwigia* sp. sect. Jussieae, interspecific hybrids can be obtained when the species used as a female has the lowest ploidy level.

Natural hybrids within section *Jussieae* have been reported between *La* (2n = 4x= 32) and *L. peploides* subsp*. stipulacea* (2n= 2x =16), with production of a triploid sterile hybrid (2n = 3x= 24) named *L. x taiwanensis* (Peng, 1990). Between *Lgg* (2n = 6x = 48) and *Lgh* (2n = 10x = 80), an octoploid hybrid was produced (2n = 8x = 64) and between *Lgg* (2n = 6x = 48) and *L. hookeri* (2n = 2x = 32), a pentaploid hybrid was produced (2n = 5x = 40) (Zardini et al., 1991; Zardini and Raven, 1992). For our *Lpm* x *Lgh* crosses, we obtained fruit production after each pollination. Despite the production of a significant seed number, very low germination was found, with no viable plants. Dandelot (2004) reported that in France, hybrids between *Lpm* and *Lgh* have never been recorded in nature, whereas hybrids have been created under experimental conditions. But if Dandelot (2004) obtained fruit from *Lpm* x *Lgh* crosses, the ability of seeds to germinate and viability of plantlets were not analyzed. As found by Dandelot, (2004), we found zero fruit production when *Lgh* was used as female.

All interspecific crosses using the lower ploidy of *Ludwigia* ssp. as female were functional and fruits were produced. But depending on the type of interspecific crosses, no viable seeds or necrotic plants were obtained. Crosses between related species or parents with different ploidy are often impossible due to post-zygotic reproductive barriers in which the hybrid progeny fails to develop or becomes sterile. Thus, in crosses between *B. napus* and a more distant species such as *Sinapis alba*, the interspecific hybridization efficiency is also extremely low and embryos need to be rescued using fertilized ovary culture (Chèvre et al., 1994). This indicated an early abortion of seeds after fertilization and the parental genome dosage in the endosperm plays an important role for seed collapse.

Interspecific hybrids between *Ludwigia* spp. in section Jussieae seem possible only if interspecific crosses occur between a female plant with lower ploidy level than male plant, and probably at a very low success rate in natura. However, observing fruit production is not enough, thus, we recommend observing seed germination, plantlet viability, plant survival, and chromosome counts.

## Conclusion

Thus, in this study we demonstrated the interest of a truly novel combination of data to identify genomic relationships and origins of polyploids in a poorly understood *Ludwigia* complex. One way to investigate phylogenetic relationship in a polyploid complex is to use flow cytometric analyses complemented with chromosome counts, as recently described for the analysis of the polyploid complex *Linum suffruticosum s.l.* (Linaceae) (Afonso et al., 2021). Another way involves (i) the use of organellar DNA (chloroplast or nuclear regions) as molecular markers as it was described for phylogenetic analysis of the genus *Isoëtes* (Pereira et al., 2019) or the diploid and autohexaploid cytotypes of *Aster amellus* (Mairal et al., 2018); or (ii) OMICS-data tools as RAD-Seq (restriction site-associated DNA sequencing) as described in the evolutionary processes of apomictic polyploid complexes on the model system *Ranunculus* (Karbstein et al., 2022). Thus, the various approaches used in this study, combining morphological and cytogenetic analyses, in situ hybridization and interspecific crosses, could constitute a first step towards phylogenetic studies of species belonging to poorly understood complexes for which there are few genomic resources.

Our results suggest that allopolyploidy played an important role in the evolutionary history of the *Ludwigia* L., section *Jussieae*, giving rise to complex relationships among species. However, some species are missing in our analyses as well as in Liu et al. (2017). The missing species of section *Jussiaea* are the four diploids, *Ludwigia peploides* (Kunth) P.H.Raven subsp. *glabrescens* (O. Kuntze) P.H.Raven, *Ludwigia peploides* subsp. *peploides*, *Ludwigia peploides* subsp. *stipulacea* (Ohwi) P.H.Raven, *Ludwigia torulosa* (Arn.) H.Hara. and the two tetraploid species, *Ludwigia hookeri* (Micheli) H.Hara, *Ludwigia peduncularis* (C.Wright ex Griseb.) M.Gómez (Hoch et al., 2015). As part of the phylogenetic relationships remains unresolved, new GISH experiments should be done with these species, especially to identify the progenitor of the unknown 2x and 4x genome of *Lgg* and *Lgh*, respectively. Furthermore, as based on morphological observations, (Zardini et al., 1991) suggested that *Lgh* may be result of interspecific hybridization between *Lgg* and *L. hookeri*, the tetraploid species

1. *L. hookeri* could be one of progenitor of missing genomes of *Lgg* and *Lgh* species.

## Acknowledgments

This research was supported by FEDER funds from Région Centre-Val de Loire and by Agence de l’eau Loire-Bretagne (grant Nature 2045, programme 9025 (AP 2015 9025). FEDER also financed the doctoral grant of L. Portillo – Lemus. Collections in Alabama were supported by the Department of Biology at the University of Alabama at Birmingham with field help from S. Heiser and S. Shainker-Connelly. The authors thank also Biogenouest (the western French network of technology core facilities in life sciences and the environment, supported by the Conseil Regional Bretagne). The authors address many thanks to PCI recommender Malika Ainouche and the two reviewers, Alex Baumel and Karol Marhold for their constructive and high-quality comments which greatly brush up the article and once again to Malika Ainouche for accepting to be recommender and organizing an efficient and constructive peer-review.

## Author Contributions

DB, LMP and OC contributed to conception and design of experiments. LPM and DB provide roots and DNA of all Lg species for *in situ* hybridization and ploidy level analysis. SAKH collected Lgg samples from USA. VH and OC acquired GISH and cytological data. BD, LMP and OC carried out analysis and interpretation of data. DB write the draft of this manuscript and DB, LPM, SAKH and OC revised the manuscript. All authors gave a final approval of the version to be published.

## Supporting Information

**Supplementary Figure S1:**
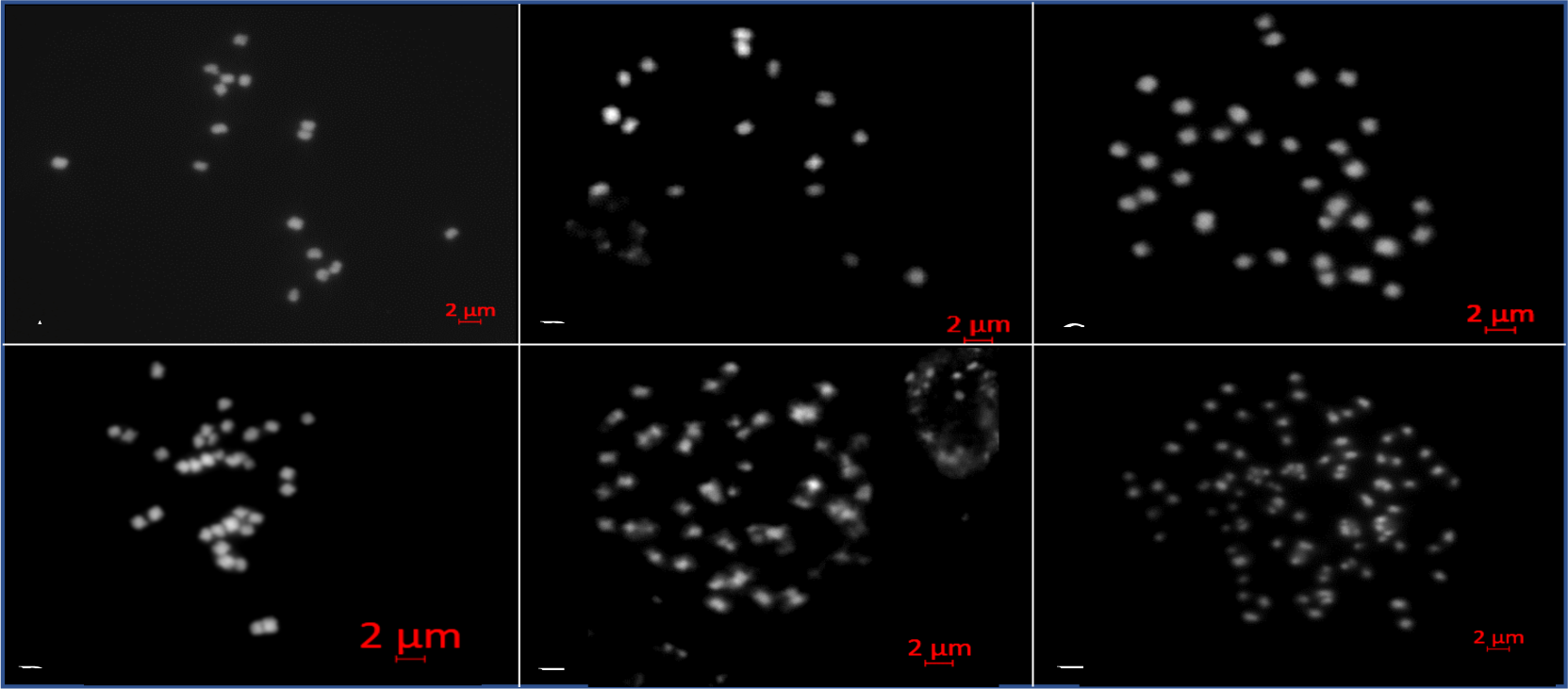
Polyploidy levels of different species of ludwigia sp section Jussieae. (A) *Ludwigia peploides* subsp. *montevidensis* chromosomes (2n=2x=16), (B) *Ludwigia helminthorrhiza* chromosomes (2n=2x=16); (C) *Ludwigia stolonifera* chromosomes (2n=4x=32); (D) *Ludwigia adscendens* chromosomes (2n=4x=32); (E) *Ludwigia grandiflora* subsp. *grandiflora* (2n=6x=48); *Ludwigia grandiflora* subsp. *hexapetala* (2n=10x=80). Chromosome number correspond to ploidy level : 16 chromosomes for diploid species (A) and (B); 32 chromosomes for tetraploid species (C) and (D); 48 chromosomes for hexaploid species (E) and 80 chromosomes for decaploid species (F).

**Supplementary Figure S2:**
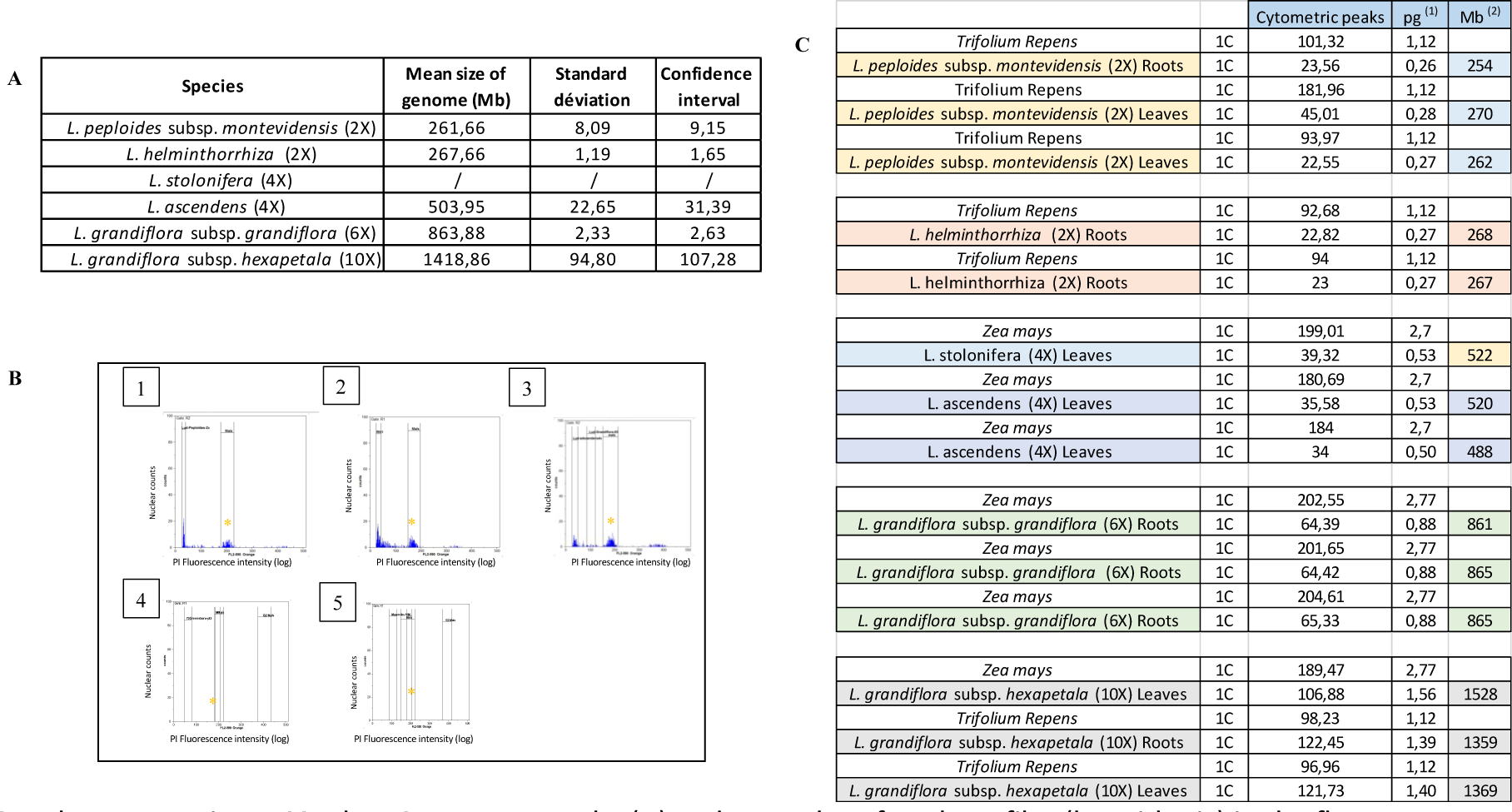
Flow Cytometry results (**A**) and examples of peak profiles (logarithmic) in the flow cytometer of nuclei stained from roots with propidium iodide (PI) (**B**). The ‘trifolium repens’ peak (1C=1,12 pg) or “Zea mays” peak (1C=2,77 pg) is used as internal standard to determinate the DNA contents of the sample nuclei (*). (1) *Ludwigia peploides* subsp. *montevidensis*; (2) *L. helminthorrhiza;* (3) *L. adscendens*; (4) *L. grandiflora* subsp. *grandiflora* and (5) *L. grandiflora* sp. *hexapetala*. ^1^ : 1 pg DNA = 978 Mbp (from Doležel et al. 2003) ; ^2^ : Zonneveld et al, 2019.

**Supplementary Figure S3:**
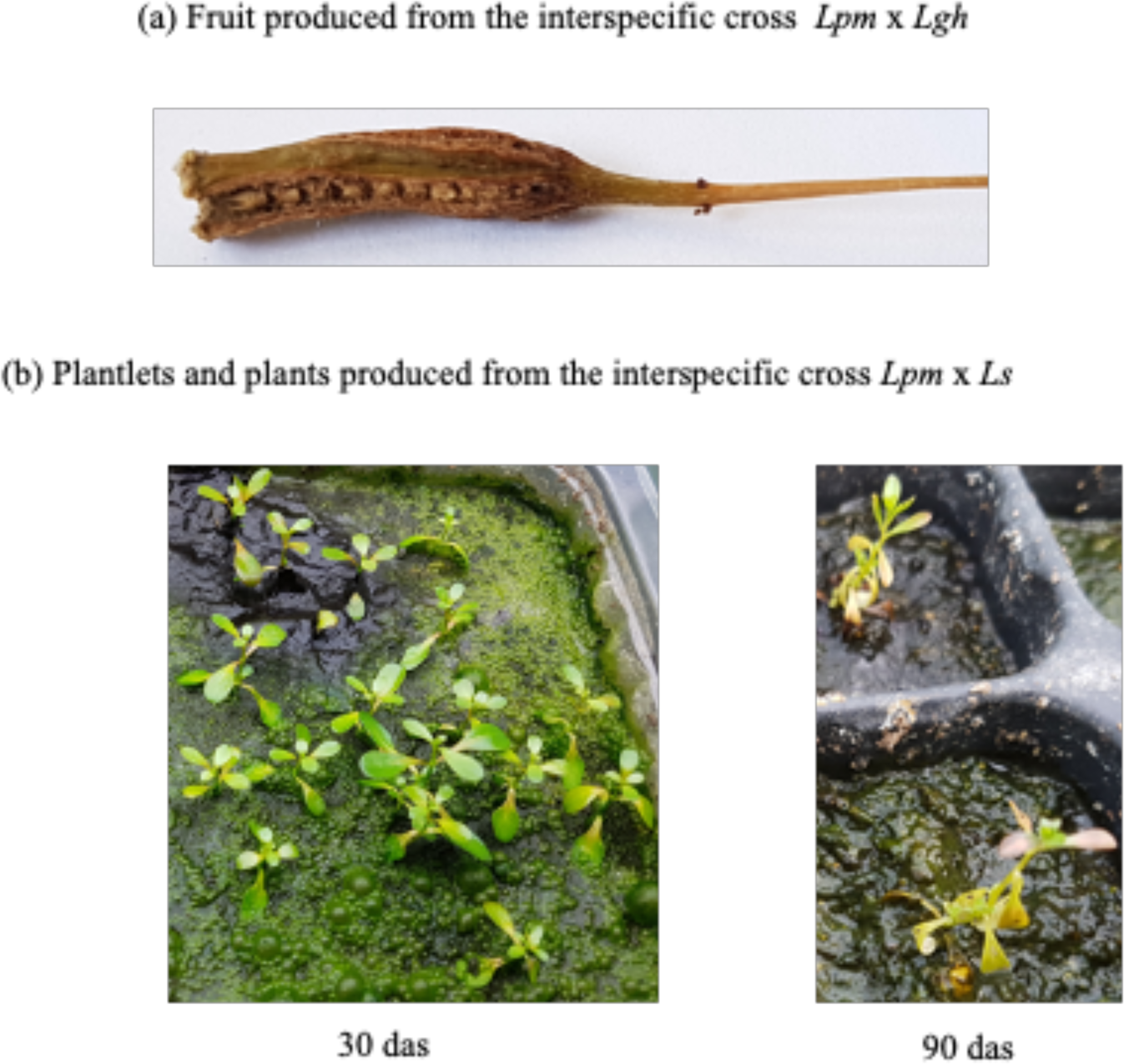
Fruit production and seedling from interspecific hybridization between *ludwigia* species possessing A genome; *Lpm* = *Ludwigia* peploides subsp. montevidensis (2n=16, AA); *Ls* = *Ludwigia stolonifera* (2n=32, AABB); *Lgh* = *Ludwigia grandiflora* subsp. *hexapetala* (2n=80, AAAABBXXXX/XXYY). (a) the seeds produced from *Lpm* x *Lgh* interspecific cross are large, which has led to the fruit bursting. (b) 30 days after seedling, green plantlets from *Lpm* x *Ls* interspecific cross were obtained. But, 60 days later, plants showed chlorotic development, stopped growing and died. Das: Number of days after seedling.

**Supplementary Figure S4:**
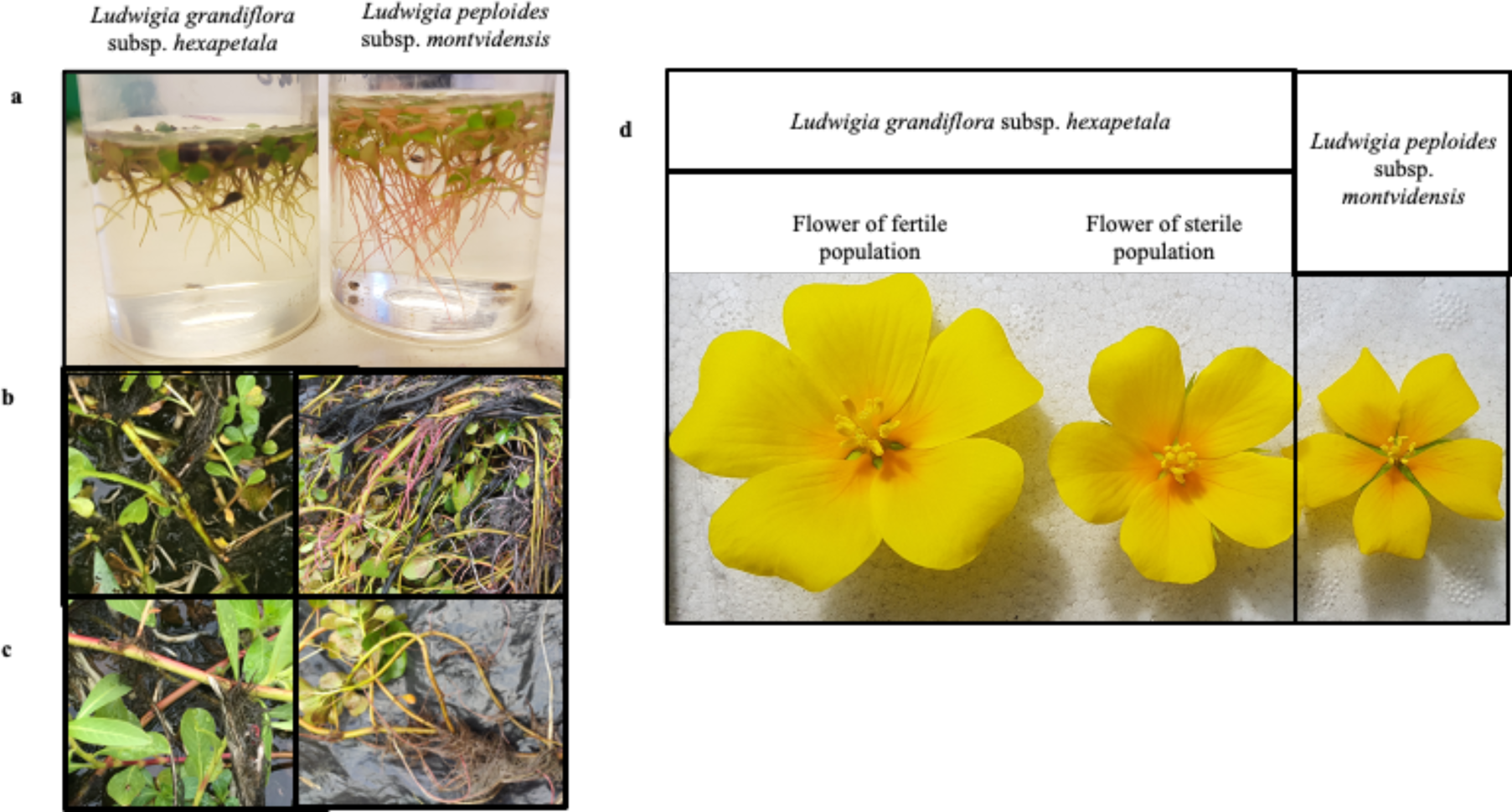
Morphological traits to distinguish *Ludwigia peploides* subsp. *montevidensis* and *Ludwigia grandiflora* subsp. *hexapetala,* (**a**) roots at seedling stage ; (**b**) adult roots in natura ; (**c**) pneumatophores in natura ; (**d**) flowers.

**Supplementary Table S1:**
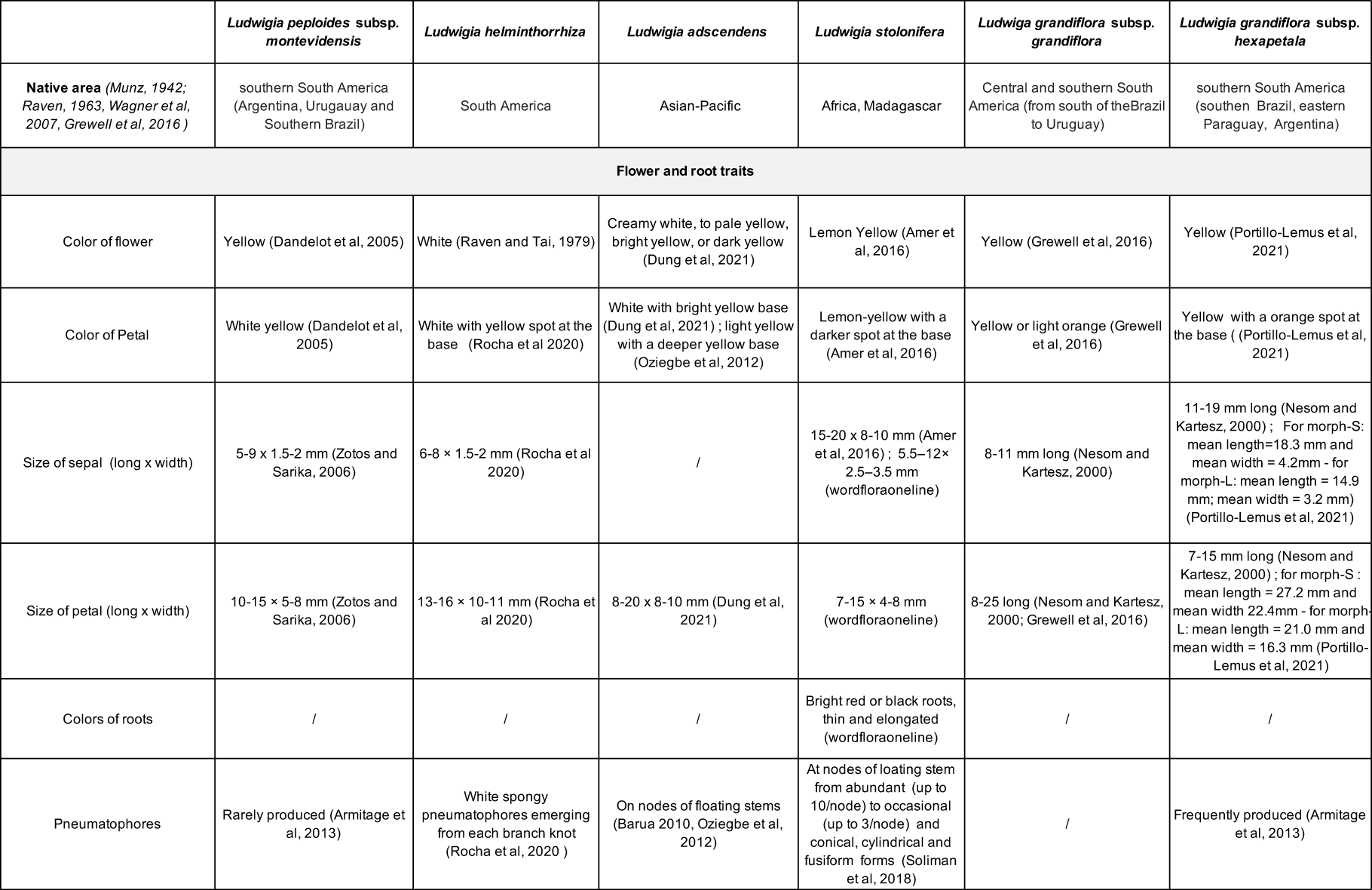
Native area and floral and root traits of the six ludwigia sp. studied. ND: not described.

|  | *Ludwigia peploides* subsp. *montevidensis* | *Ludwigia helminthorrhiza* | *Ludwigia adscendens* | *Ludwigia stolonifera* | *Ludwigia grandiflora* subsp. *grandiflora* | *Ludwigia grandiflora* subsp. *hexapetala* |
| --- | --- | --- | --- | --- | --- | --- |
| **Native area** (Munz, 1942; Raven, 1963, Wagner et al, 2007, Grewell et al, 2016 ) | southern South America (Argentina, Uruguay and Southern Brazil) | South America | Asian-Pacific | Africa, Madagascar | Central and southern South America (from south of the Brazil to Uruguay) | southern South America (southern Brazil, eastern Paraguay, Argentina) |
| **Flower and root traits** |  |  |  |  |  |  |
| Color of flower | Yellow (Dandelot et al, 2005) | White (Raven and Tai, 1979) | Creamy white, to pale yellow, bright yellow, or dark yellow (Dung et al, 2021) | Lemon Yellow (Amer et al, 2016) | Yellow (Grewell et al, 2016) | Yellow (Portillo-Lemus et al, 2021) |
| Color of Petal | White yellow (Dandelot et al, 2005) | White with yellow spot at the base (Rocha et al 2020) | White with bright yellow base (Dung et al, 2021) ; light yellow with a deeper yellow base (Oziegbe et al, 2012) | Lemon-yellow with a darker spot at the base (Amer et al, 2016) | Yellow or light orange (Grewell et al, 2016) | Yellow with a orange spot at the base ((Portillo-Lemus et al, 2021) |
| Size of sepal (long x width) | 5-9 x 1.5-2 mm (Zotos and Sarika, 2006) | 6-8 x 1.5-2 mm (Rocha et al 2020) | / | 15-20 x 8-10 mm (Amer et al, 2016) ; 5.5-12x 2.5-3.5 mm (wordfloraonline) | 8-11 mm long (Nesom and Kartesz, 2000) | 11-19 mm long (Nesom and Kartesz, 2000) ; For morph-S: mean length=18.3 mm and mean width = 4.2mm - for morph-L: mean length = 14.9 mm; mean width = 3.2 mm) (Portillo-Lemus et al, 2021) |
| Size of petal (long x width) | 10-15 x 5-8 mm (Zotos and Sarika, 2006) | 13-16 x 10-11 mm (Rocha et al 2020) | 8-20 x 8-10 mm (Dung et al, 2021) | 7-15 x 4-8 mm (wordfloraonline) | 8-25 long (Nesom and Kartesz, 2000; Grewell et al, 2016) | 7-15 mm long (Nesom and Kartesz, 2000) ; for morph-S : mean length = 27.2 mm and mean width 22.4mm - for morph-L: mean length = 21.0 mm and mean width = 16.3 mm (Portillo-Lemus et al, 2021) |
| Colors of roots | / | / | / | Bright red or black roots, thin and elongated (wordfloraonline) | / | / |
| Pneumatophores | Rarely produced (Armitage et al, 2013) | White spongy pneumatophores emerging from each branch knot (Rocha et al, 2020 ) | On nodes of floating stems (Barua 2010, Oziegbe et al, 2012) | At nodes of floating stem from abundant (up to 10/node) to occasional (up to 3/node) and conical, cylindrical and fusiform forms (Soliman et al, 2018) | / | Frequently produced (Armitage et al, 2013) |

**Supplementary Table S2:** Reproductive success after controlled interspecific crosses between different *Ludwigia* L. spp. belonging to the section *Jussiaea*. Interspecific hybridization (female x male) between the three species, *Ludwigia peploides* subsp. *montevidensis* (Lpm), *Ludwigia stolonifera* (Ls) and/or *Ludwigia grandiflora* subsp. *hexapetala* (Lgh, AAAA BB XXXX/XXYY) used as female or male. All species possess same genome A: Lpm (2x, AA); Ls (4x, AABB); Lgh (10x, AAAA BB XXXX or XXYY). Number of plantlets and plants were counted three (21 days) and 8 weeks (56 days) after seed germination, respectively. NA: data not available. (+/-= confidence interval, lll=0.05). For control interspecific crosses *Lgh* x *Lgh* and *Lpm* x *Lpm*, a set of randomly selected plantlets were followed until 56 days after seed germination.

| Controlled interspecific crosses | *Lpm* x *Ls* | *Lpm* x *Lgh* | *Ls* x *Lpm* | *Ls* x *Lgh* | *Lgh* x *Lpm* | *Lgh* x *Ls* | *Lgh* x *Lgh* | *Lpm* x *Lpm* |
| --- | --- | --- | --- | --- | --- | --- | --- | --- |
| Number of cross pollination | 8 | 25 | 10 | 2 | 10 | 10 | 75 | 45 |
| Number of fruits | 8 | 25 | 0 | 2 | 0 | 0 | 75 | 45 |
| Mean length of fruit (mm) | 15.08 (+/- 0.78) | 16.64 (+/- 0.82) | / | NA | / | / | 7 | NA |
| Mean fruit weight (g) | 62.04 (+/- 6.46) | 64.64 (+/- 6.02) | / | NA | / | / | NA | NA |
| Number of total seed | 221 | 1101 | / | 47 | / | / | 3750 | 1980 |
| Number of germinated seeds | 118 | 34 | / | 0 | / | / | 3375 | 1881 |
| Number of plantlets 21 days | 118 | 3 | / | 0 | / | / | 3750 | 1881 |
| Number of plants 56 days | 0 | 0 | / | 0 | / | / | 100 from a set of 100 | 50 from a set of 50 |

